# A Multi-pronged Screening Approach Targeting Inhibition of ETV6 PNT Domain Polymerization

**DOI:** 10.1101/2020.08.20.258525

**Authors:** Chloe A. N. Gerak, Si Miao Zhang, Aruna D. Balgi, Ivan J. Sadowski, Richard B. Sessions, Lawrence P. McIntosh, Michel Roberge

**Author notes:** **Corresponding Author** Michel Roberge, Department of Biochemistry and Molecular Biology, University of British Columbia, Vancouver, BC, Canada.

## Abstract

ETV6 is an ETS family transcriptional repressor for which head-to-tail polymerization of its PNT domain facilitates cooperative binding to DNA by its ETS domain. Chromosomal translocations frequently fuse the ETV6 PNT domain to one of several protein tyrosine kinases. The resulting chimeric oncoproteins undergo ligand-independent self-association, autophosphorylation, and aberrant stimulation of downstream signaling pathways leading to a variety of cancers. Currently, no small molecules inhibitors of ETV6 PNT domain polymerization are known and no assays targeting PNT domain polymerization have been described. In this study, we developed complementary experimental and computational approaches for identifying such inhibitory compounds. One mammalian cellular approach utilized a mutant PNT domain heterodimer system covalently attached to split *Gaussia* luciferase fragments. In this protein fragment complementation assay, inhibition of PNT domain heterodimerization reduces luminescence. A yeast assay took advantage of activation of the reporter *HIS3* gene upon heterodimerization of mutant PNT domains fused to DNA-binding and transactivation domains. In this two-hybrid screen, inhibition of PNT domain heterodimerization prevents cell growth in medium lacking histidine. The Bristol University Docking Engine (BUDE) was used to identify virtual ligands from the ZINC8 library predicted to bind the PNT domain polymerization interfaces. Over 75 hits from these three assays were tested by NMR spectroscopy for binding to the purified ETV6 PNT domain. Although none were found to bind, lessons learned from this study may facilitate future approaches for developing therapeutics that act against ETV6 oncoproteins by disrupting PNT domain polymerization.

## Introduction

The *ETV6* gene, known also as *TEL* (<u>t</u>ranslocation <u>E</u>TS <u>l</u>eukemia), encodes an ETS family transcriptional repressor with roles in embryonic development and hematopoietic regulation.^1,2,3^ This gene is frequently rearranged in chromosomal translocations to form fusion oncoproteins linked with various cancers including leukemias, lymphomas, carcinomas, and sarcomas.^4–6^ Over 40 different such translocations are known to exist, including those leading to the *ETV6-PDGFRB, ETV6-NTRK3, ETV6-ABL1/2,* and *ETV6-JAK2* gene fusions.^4,6–9^

ETV6 is a modular protein composed of an N-terminal self-associating PNT (or SAM) domain, a disordered central region reported to interact with co-repressors, and a C-terminal DNA-binding ETS domain.^10,11^ Most of the oncogenic *ETV6* translocations lead to chimeras with the PNT domain fused to the protein tyrosine kinase (PTK) domain from a receptor tyrosine kinase. Crucial to their oncogenic properties is the propensity of the ETV6 PNT domain to tightly self-associate into an open-ended, left-handed helical polymer via head-to-tail binding of two relatively flat, hydrophobic interfaces.^12–14^ PNT domain polymerization enables ligand-independent autophosphorylation and activation of the PTK domain. This stimulates downstream cellular pathways, such as the PI3K/Akt and Ras-MAPK signaling cascades, and ultimately causes cellular transformation.^15–17^

One well-characterized *ETV6* translocation encodes the PNT domain fused to the PTK domain of neurotrophin tyrosine receptor kinase-3 (NTRK3). The resulting protein, named EN (ETV6-NTRK3), displays oncogenic properties including phenotypic transformation and soft agar colony formation of several experimental cell lines, as well as tumor formation in nude mice.^6,17^ The ETV6 PNT domain polymerization interfaces, called the mid-loop (ML) and end-helix (EH) surfaces, have hydrophobic cores surrounded by charged residues.^18^ The mutation of one of two key hydrophobic residues to a charged residue (A93D on the ML surface, or V112E or V112R on the EH surface, according to the human ETV6 numbering) disrupts polymerization.^14,18^ Introduction of these mutations into *EN* expressing cell lines prevents EN polymerization, PTK activation and cellular transformation.^13^ Similarly, mutation of the K99-D101 charge pair bridging the PNT domain interfaces weakens polymerization and inhibits the transformation activity of EN in NIH3T3 cells.^19^ Co-expression of an isolated PNT domain also has a dominant negative effect, preventing cellular transformation.^13^ Together, these studies indicate inhibition of PNT domain polymerization as a viable therapeutic strategy against cancers driven by *ETV6* chromosomal translocations.

We hypothesized that small molecules that prevent the self-association, and hence oncogenic properties, of ETV6 chimeras containing the PNT domain might serve as a potential broad spectrum therapy against many ETV6 PNT domain-containing oncoproteins and avoid toxicities associated with perturbing the normal activities of receptor PTKs. Furthermore, although approximately one-third of the 28 ETS transcription factor family members in humans possess a PNT domain, only ETV6 and likely ETV7 polymerize.^20–22^ Thus a molecule that selectively inhibits ETV6 PNT domain polymerization may have few side effects.

Protein-protein interactions (PPIs) are challenging to disrupt, and hence we undertook complementary cellular and virtual screening strategies in the attempt to discover inhibitors of ETV6 PNT domain polymerization. Central to our approach is the use of PNT domains with monomerizing mutations in the EH or ML surfaces that can still associate with low nM affinity through their remaining complementary wild-type interfaces.^14,18^ The soluble “heterodimer” serves as a model of the insoluble polymer, opening the door for *in vitro* and *in vivo* screens. In brief, both a mammalian cell-based assay utilizing a protein-fragment complementation approach with split *Gaussia* luciferase^23^ (Fig. 1A) and a yeast two-hybrid assay^24^ (Fig. 1B) were developed and used to screen chemical libraries for potential inhibitors. In parallel, large scale virtual screening using the Bristol University Docking Engine (BUDE) was performed to identify theoretical compounds that might bind to the PNT domain polymerization interfaces.^25,26^ Subsequently, candidate compounds were tested for inhibitory effects in cellular assays and for binding to the isolated ETV6 PNT domain as monitored by nuclear magnetic resonance (NMR) spectroscopy. Although no inhibitory compounds were successfully identified, the development, validation and implementation of these assays will be discussed herein.

**Figure 1.**
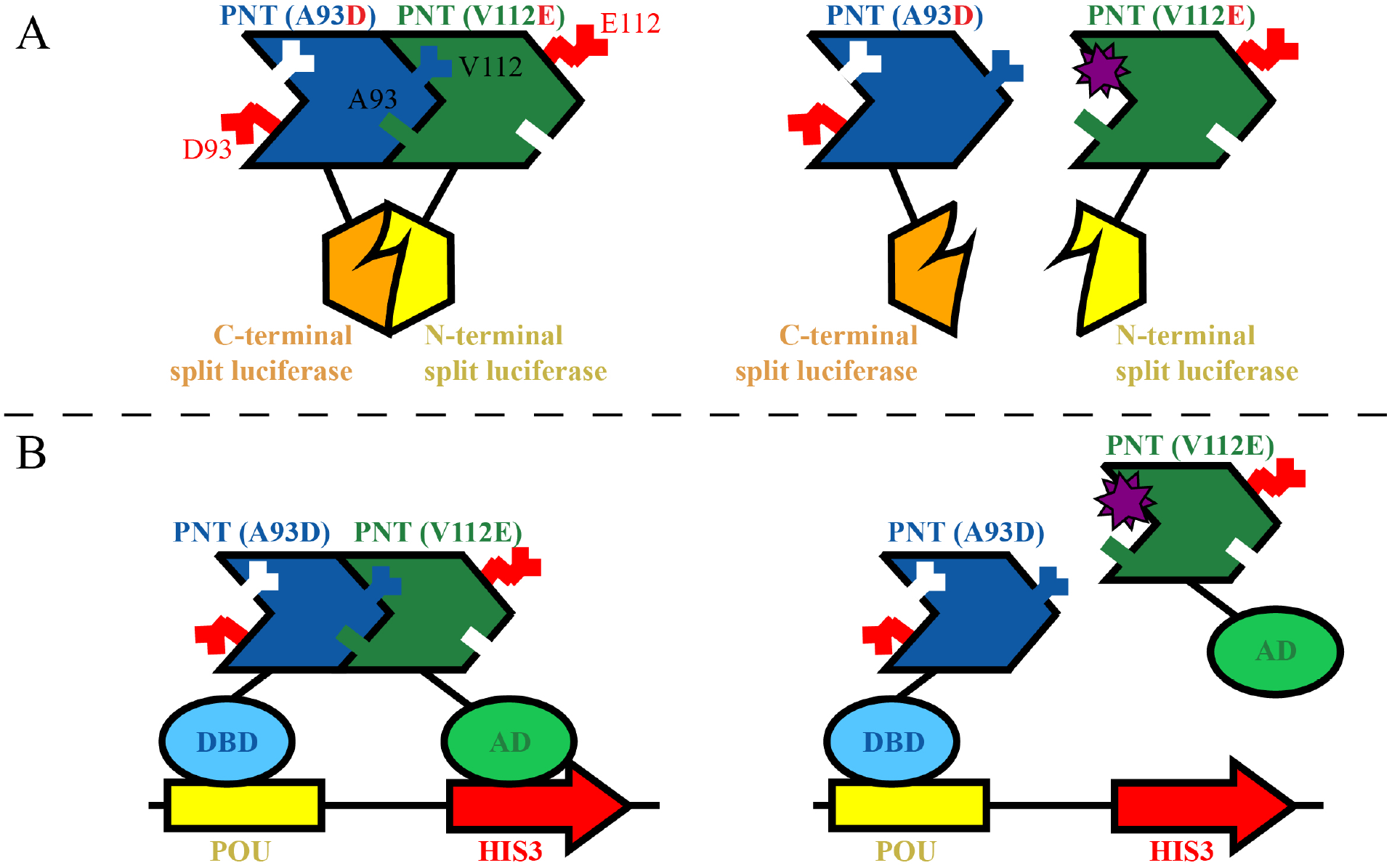
(**A**) The general principle behind the split-luciferase PCA. When the A93D- and V112E- PNT domains heterodimerize (left), the split luciferase fragments assemble into an active enzyme which, upon addition of substrate, generates a luminescence output. If a small molecule (star) disrupts the interaction (right), either by directly binding the interface or by inducing an allosteric conformation change, then the equilibrium shifts away from reconstitution of the luciferase fragments, resulting in reduced luminescence. (**B**) In the yeast two-hybrid approach, the PNT domains are covalently linked to either a *POU* DNA-binding domain (DBD) or an activating domain (AD). Heterodimerization induces expression of the *HIS3* gene to allow cell growth in medium lacking His (left). If a small molecule disrupts the interaction (right), there is no expression of the *HIS3* gene and no yeast growth. Addition of His to the medium enables yeast growth in the absence or presence of an inhibitor of the PNT domain interaction or the histidine biosynthetic pathway. These cartoons are highly schematic, especially with respect to the folding of the luciferase fragments.

## Materials and Methods

### Chemicals for Screening

Screening compounds for the cellular assays consisted of 16,000 compounds from the Maybridge Hitfinder collection, 10,000 compounds from the ChemBridge DIVERset collection, 1,120 compounds from Prestwick Chemicals, 1,280 compounds from the Sigma LOPAC library, 2,000 compounds from the Microsource Spectrum collection, 2,697 compounds from the Selleck L1700 Bioactive Compound library, and 500 compounds from Biomol. The compounds were stored in 96-well plates at −25 °C as 5 mM stock solutions in DMSO. In addition, a small molecule library targeting PPIs consisting of 1,534 compounds was provided by the Perturbation of Protein-Protein Interactions (PoPPI) collaborative program (UK). Candidate compounds from the BUDE screening assay were purchased from MolPort for testing in both cellular assays and by NMR spectroscopy.

### Vectors and Cloning for the Split Luciferase Protein-fragment Complementation Assay

The split luciferase protein-fragment complementation assay (PCA) was based on the protocol of Michnick and co-workers.^23^ Sequences encoding ETV6^43-125^ (residues 43-125 of human ETV6, encompassing the PNT domain; Genbank Gene ID: 2120) with either an A93D or V112E substitution were cloned into either the modified mammalian expression vector pcDNA3.1/Zeo(+) or pcDNA3.1/Neo(+) at the 5’-end of sequences for humanized Gaussian Luciferase (hGLuc) fragments (Supplemental Table S1). The resulting constructs encoded either A93D- or V112E-PNT domain, a (GGGGS)_2_ flexible linker and either hGLuc(1)^1-93^ or hGLuc(2)^94-196^. The latter are described herein as N-Luc or C-Luc, respectively. A control set of plasmids containing leucine zippers as the dimerization domains were also provided by Dr. Michnick.^23^

### Mammalian Cell Culture

Human embryonic kidney cells 293 (HEK293) cells were cultured in Dulbecco’s Modified Eagle Medium (DMEM; Gibco) supplemented with 10% fetal bovine serum (FBS; Sigma) and 1% antibiotic-antimycotic (Gibco). Unless otherwise noted, this medium was used in all experiments. Stably expressing transformants were treated additionally with either or both 400 μg/mL G418 (Gibco) and 50 μg/mL zeocin (Invitrogen). Cells were incubated at 37 °C with 5% CO_2_, and passaged upon reaching approximately 75-85% confluency.

### Transient Expression for Validation of the PCA

HEK293 cells were seeded in 96-well clear bottom black microplates (Corning #6005182) at 15,000 cells/well and incubated at 37 °C. After 24 h, the medium was aspirated and 100 μL of fresh medium added. For transfection of a single species of DNA, 20 ng/μL of plasmid was added to OPTIMEM (Gibco), and for transfection of two species of DNA, 10 ng/μL of each plasmid was added to OPTIMEM. Lipofectamine 2000 (Invitrogen) was diluted to 8% in OPTIMEM and added to the DNA at a 1:1 v:v ratio and incubated for 5 min at room temperature. After incubation, 10 μL of the total prepared DNA, OPTIMEM and Lipofectamine 2000 were added to each well. Cells were incubated at 37 °C for either 24, 48 or 72 h. Prior to the luminescence reading, 50 μL of cell medium was removed and an equal volume of NanoFuel GLOW Assay (Nanolight Technology) for *Gaussia* luciferase was then added to each well. After incubation in the dark at ambient temperature for 5 min, the luminescence output was read for one second with a Varioskan LUX multimode microplate reader.

### Establishment of a Stably Expressing PNT Domain PCA System

HEK293 cells were seeded in 6-well microplates at 400,000 cells/well and grown overnight to approximately 80% confluence. Cell medium was changed and the cells were transfected with Lipofectamine 2000 utilizing 25 ng/μL DNA for single plasmid transfections or 12.5 ng/μL DNA for each plasmid in a double plasmid transfection. After 24 h, the medium was aspirated, fresh medium was added and the cells were incubated again overnight. After 48 h, selection was introduced by incubating cells with medium supplemented with the corresponding antibiotic(s), replacing medium every 2-3 days, and splitting cells when 80% confluency was achieved. The resulting stable transformants were stored in liquid nitrogen.

### High-throughput PCA Screening

For screening, stable transformants of A93D-PNT/N-Luc(Neo) and V112E-PNT/C-Luc(Zeo) were plated at 40,000 cells/well in 96-well clear bottom black microplates (Corning #6005182) and incubated overnight at 37 °C. Compound plates were thawed and compounds were added to cells using a BioRobotics BioGrid Robot Microarrayer Model equipped with a 96-pin tool with either 0.7 mm or 0.4 mm diameter pins. After a 4 h incubation at 37 °C, a 1:1 ratio of NanoFuel GLOW Assay reagent was added to the cells and plates were incubated in the dark at ambient room temperature for 15 min. Luminescence output was then read with a Varioskan LUX multimode or BioTek Neo 2 microplate reader.

### Secondary PCA

Compounds that yielded a lower luminescence in the initial PCA screens were re-tested at different concentrations against cells stably expressing the split luciferases fused to ETV6 PNT domains or leucine zippers. The latter served as a specificity control. Cells were seeded at 40,000 cells per well in 96-well plates and incubated at 37 °C overnight. The selected compounds were added to wells in duplicate at final concentrations of 1, 3, 10 and 30 μM. After 4 h incubation, cells were visually examined to determine if the compound caused toxicity or precipitated from solution. Luminescence was read as previously described.

### Yeast Two-hybrid Assay Development

The two-hybrid assay consists of bait and prey plasmids and a yeast reporter strain. The bait plasmids expressed, from a constitutive *ADH1* promoter, the Oct1 POU DNA-binding domain alone (pIS341) or a fusion between the Oct1 POU DNA-binding domain and the A93D-PNT domain (pIS586). They are ARS-CEN (single copy) plasmids with a *TRP1* marker. The prey plasmids expressed the NLS-B42 activation domain (AD) alone (pIS580) or a fusion between the AD and V112E-PNT domain (pIS591). These genes are under control of an inducible *GAL1* promoter on 2 micron (multicopy) plasmids with a *LEU2* marker.

*Saccharomyces cerevisiae* strain ISY361 was constructed from strain W303. It contains two integrated reporter genes. The *HIS3* reporter with a minimal core promoter was integrated at an *ade8* disruption with plasmid pIS452.^27^ The *lacZ* reporter was integrated at a *lys2* disruption with pIS341.^27^ The expression of the two reporter genes is controlled by 4 upstream POU binding sites. The genotype is *MATα*, *ade2-*1, *his3-11,15*, *leu2-3,112*, *trp1-1*, *ura3-1*, *can1-100*, *lys2::POU ops-LACZ*, *ade8::POU ops-HIS3.*

ISY361 was transformed with bait plasmid pIS586 and prey plasmid pIS591 to generate the strain ISY361+/+ expressing POU-A93D bait and AD-V112E prey. A strain containing bait plasmid pIS586 and prey plasmid pIS580 lacking the V112E domain was also generated to serve as a control and is referred to as ISY361+/−.

### High-throughput Yeast Two-hybrid Screening

Yeast media were prepared with reagents obtained from Becton Dickinson and Sunrise Science Products. Strains were grown overnight at 30 °C with agitation in Synthetic Complete (SC) medium lacking Leu and Trp and containing 2% glucose. Cells were harvested by centrifugation at 4,700 g for 5 min, pellets were rinsed twice with sterile distilled water and cells were suspended at OD_595_ 0.01 in SC medium lacking Leu, Trp and His and containing 2% galactose instead of glucose. The suspension also contained 2 mM 3-amino-1,2,4-triazole, which is used to minimize the effect of basal expression of *HIS3*.^28,29^ Cell suspension (100 μL) was distributed to wells of sterile clear flat bottom polystyrene 96-well microplates (Costar # 3370) using a dispensing 8-channel pipettor. Eight wells were reserved for blanks. Chemicals were added to each well using a Biorobotics Biogrid II robot equipped with either a 0.7 mm or 0.4 mm diameter 96-pin tool. The yeast plates were incubated at 30 °C in a humidified chamber without agitation for 48 h. The cells were suspended by gently vortexing for 1 min and OD_595_ readings were obtained using an Opsys MR 96-well plate reader (Dynex Technologies). OD_595_ readings of wells containing only medium were defined as 0% growth and OD_595_ readings of wells containing yeast but no screening chemicals were defined as 100% growth.

The compounds were tested at either a final concentration of 10 or 15 μM. Compounds showing growth inhibition were typically re-tested at two different concentrations in duplicate in medium lacking His and in medium containing 20 μg/mL L-His. Compounds showing more growth inhibition in medium lacking His than in medium containing His were re-tested in two or more replicates over a concentration range.

### BUDE Virtual Screening

Virtual ligand screening was carried out on the University of Bristol’s high performance computing system BlueCrystal with the docking program BUDE (Version 1.2.9)^25^ utilizing the University of California San Francisco ZINC8 virtual ligand database.^30^ Coordinates of monomer subunits were taken from the X-ray crystallographic structure of the polymeric ETV6T PNT domain (PDB: 1LKY) and used as representative of the wild-type interfaces in the A93D-PNT or V112R-PNT domain backgrounds. In brief, the protein structure, known as the receptor, was placed as a mol2 file with the origin at the wild-type A93 or V112 residue for the V112R- or A93D-PNT domain, respectively. Centered on the origin, the docking grid search volume was a 15 Å × 15 Å × 15 Å cube. Members of the ZINC8 library, consisting of greater than 8 million ligands (each having approximately 20 conformers per ligand generated), were tested for docking around the origin.

A second BUDE screen with residues of the intermolecular K99-D101 salt bridge set as the origins. In addition, an ensemble of 10 different structures, obtained from 10 ns steps of a 100 ns molecular dynamic (MD) simulation performed with GROMACS,^31^ were used as the receptors. Docking was carried out using the top 200,000 compounds, and their conformers, that exhibited the lowest binding energies in the first BUDE screen, described above.

### Molecular Dynamics Simulations on Docked Candidates

Ligands that were targeted for the interfacial residues of the ETV6 PNT domain were ranked on their predicted free energy of binding. The top 500 compounds to each interface underwent a short 10 ns MD simulation of the ligand-protein complex to determine if they maintained a stable interaction. In brief the MD simulations were performed with AMBER (Version 16)^32^ using the FF14SB-ildn forcefield, TIP3P water and ligand parameters taken from the GAFF (General Amber Force Field).^33^ The full 10 ns simulations were run with 2 fs integration step size while maintaining a temperature of 300 K and pressure of 1 bar. Resulting ligand root-mean-squared deviation (RMSD) time courses were calculated for the trajectories relative to the initial, midpoint and final poses using CPPTRAJ.^34^ In addition, the trajectories were visualized using VMD (Version 1.9.2) software.^35^

### BUDE Candidate Selection and Testing

Compounds identified by virtual docking underwent several iterations of selection. First, the top ranked poses with the lowest calculated binding energies were manually inspected in Chimera (Version 1.13.1).^36^ For the ligands targeted to the interface, those that had duplicate conformers with low RMSDs during 10 ns MD simulations were preferentially selected over those with high RMSDs or that dissociated. The generated list of potential compounds was further refined by considering their commercial availability and selected to give a range of chemical diversity targeting the EH or ML surfaces of the ETV6 PNT domain. In total, 50 compounds were purchased to target the core interfacial residues. Of these, 16 targeted the ML surface around A93 in the V112E-PNT domain structure, 32 targeted the EH surface around V112 in the A93D-PNT domain structure, and 2 targeted both. In addition, 10 compounds were purchased to target the K99-D101 salt bridge (6 targeted D101 and 4 targeted K99).

### Testing of Compound Binding by NMR Spectroscopy

Candidate compounds were tested *in vitro* for binding to the ^15^N-labeled PNT domain via ^15^N-HSQC monitored titrations recorded at 25 °C with Bruker Avance 500 or 600 MHz spectrometers. Isotopically labeled ETV6^43-125^ with either an A93D or V112E substitution was expressed in *Escherichia coli* and purified as described previously.^19^ Purchased compounds were dissolved to 50 mM stock solutions in DMSO. Purified protein samples were at a final concentration of 150 μM and volume of 450 μL in a standard buffer (20 mM MOPS, 50 mM NaCl, and 0.5 mM EDTA at pH 7.0 for A93D-PNT domain and pH 8.0 for V112E-PNT domain) with D_2_O (5% v/v) added for signal locking. All compounds were tested at a minimum of 2:1 molar ratio a maximum of 20:1 molar ratio compound to protein. Control titrations with DMSO were also recorded.

## Results

### Development of a PCA to Screen for Inhibitors of PNT Domain Association

Initially, we established and characterized a PCA for monitoring heterodimerization of the A93D- and V112E-PNT domains based on the split *Gaussia* luciferase methodology.^23^ Various fusion proteins containing the mutant ETV6 PNT domains and either N-Luc or C-Luc fragments were transiently expressed in HEK293 cells (Supplemental Table S2). Cell culture medium alone or cells exposed to the reagents needed for the transient transfections showed luminescence readings of ~ 20 on a Varioskan LUX multimode microplate reader. When expressed alone, V112E-PNT/C-Luc, A93D-PNT/C-Luc and A93D-PNT/N-Luc also produced relatively low luminescence readings of ~ 600, either 48 or 72 h post-transfection. Co-expressing A93D- and V112E-PNT domains that can heterodimerize, but fused to the same luciferase fragment, also resulted in a low luminescence reading of ~ 70. Introduction of the complementary N-Luc and C-Luc luciferase fragments, each linked to the A93D-PNT domain yielded a higher luminescence of 2500. However, this was still low when compared to combinations of A93D-PNT/C-Luc and V112E-PNT/N-Luc or A93D-PNT/N-Luc and V112E-PNT/C-Luc, that showed readings 48 h post-transfection of 320,000 and 91,000, respectively. The luminescence readings of these combinations diminished 72 h post-transfection. The higher luminescence readings of the A93D-PNT/C-Luc and V112E-PNT/N-Luc combination compared with the reciprocal combination may indicate that one orientation of the luciferase fragments is more favorable to reconstitution of the active enzyme than the other. Based on these initial studies, the constructs encoding A93D-PNT/C-Luc and V112E-PNT/N-Luc were expressed stably in HEK293 cells to facilitate high-throughput screening.

There are no known inhibitors of ETV6 PNT domain polymerization to use as controls. However, the isolated PNT domain has a dominant-negative effect on cells expressing EN, reverting morphology back to wild-type.^19^ Thus, we transiently transfected the A93D-PNT domain into the stably expressing HEK293 cells to test for an expected reduction in the luminescence reading due to competition with the A93D-PNT/N-Luc for the V112E-PNT/C-Luc binding interface (Fig. 2A). At 24 h post-transfection, the A93D-PNT domain caused a significant decrease in luminescence with this cell line, but not with control cells expressing luciferase fragments fused with leucine zippers. At 48 h post-transfection, both systems were affected, but the decrease was most pronounced with the cells for the PNT domain PCA. To a much lesser extent, a transiently transfected empty pcDNA3.1 vector also reduced luminescence for the two systems 48 h post-transfection. Collectively, these controls define the expected sensitivity to inhibition of PNT domain heterodimerization.

**Figure 2.**
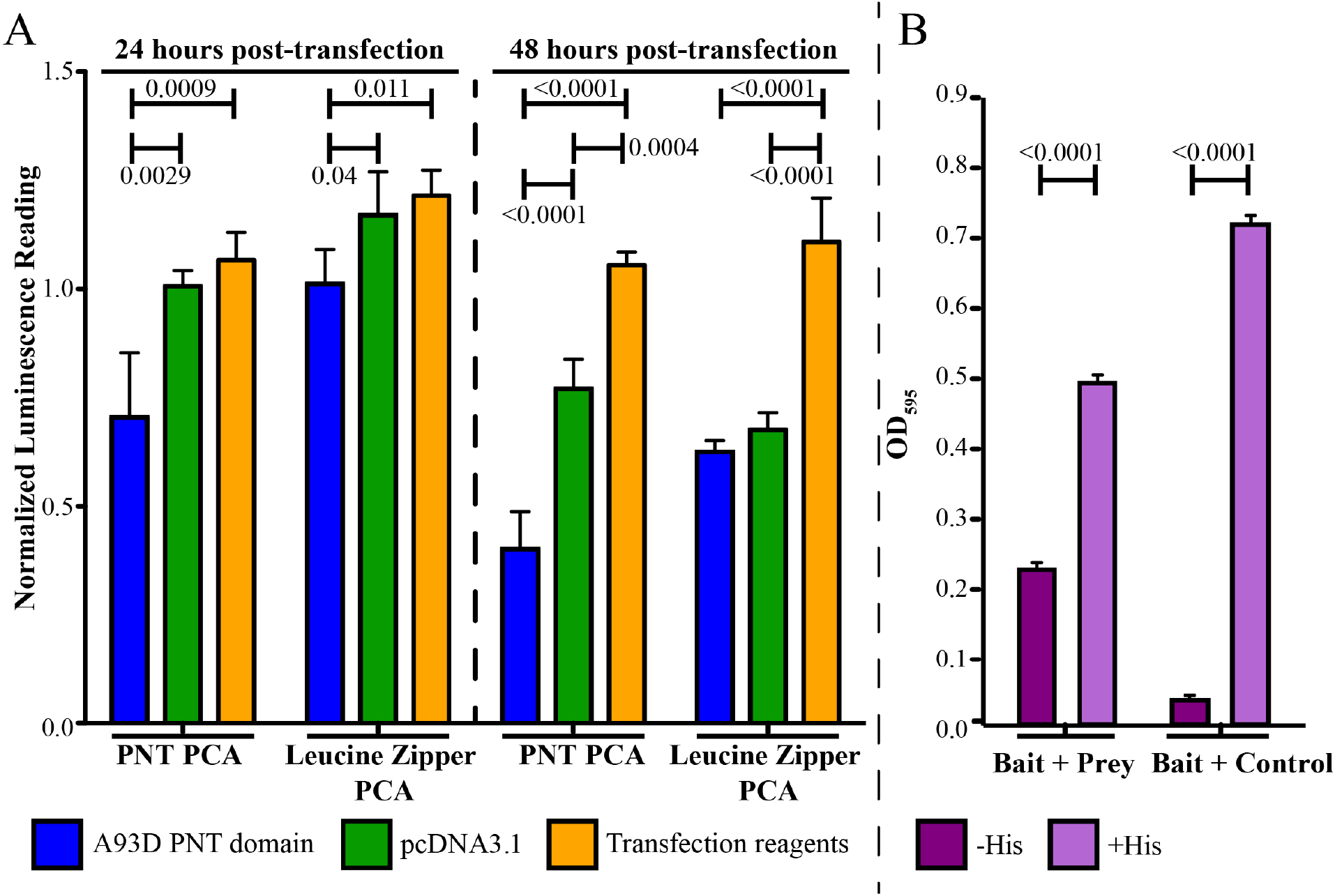
(**A**) The A93D-PNT domain-expressing and empty pcDNA3.1 vectors were transiently transfected into the PCA systems with luciferase fragments fused to either complementary PNT domains or leucine zippers. Controls with only transfection reagents were also run in parallel. Luminescence readings were normalized to untreated PNT domain or leucine zipper PCAs cells, respectively, with means and standard deviations of the 3 replicates. One-way ANOVA statistical analyses were performed and the calculated P values are given above the horizontal bars. (B) Yeast growth as detected by OD_595_. Co-expression of the bait (A93D-PNT Domain/POU DBD) + prey (V112E-PNT domain/AD) plasmids enables ISY361+/+ yeast growth with (light purple) and without (dark purple) addition of His. In contrast, without added His, no growth was seen for ISY361+/− yeast containing the bait plasmid and an empty control plasmid that lacks the complementary PNT domain. Each condition was replicated 5 times.

### High-throughput Screening using the PNT Domain PCA

In total, ~ 18,000 compounds were screened with the PNT domain PCA assay. Plates were analyzed individually due to the limited stability of the luminescence signal over time and compounds of interest were identified as having a luminescence reading that was more than two standard deviations away from the average luminescence reading of a plate (Fig. 3A). An overall ~ 1 % hit rate was observed.

**Figure 3.**
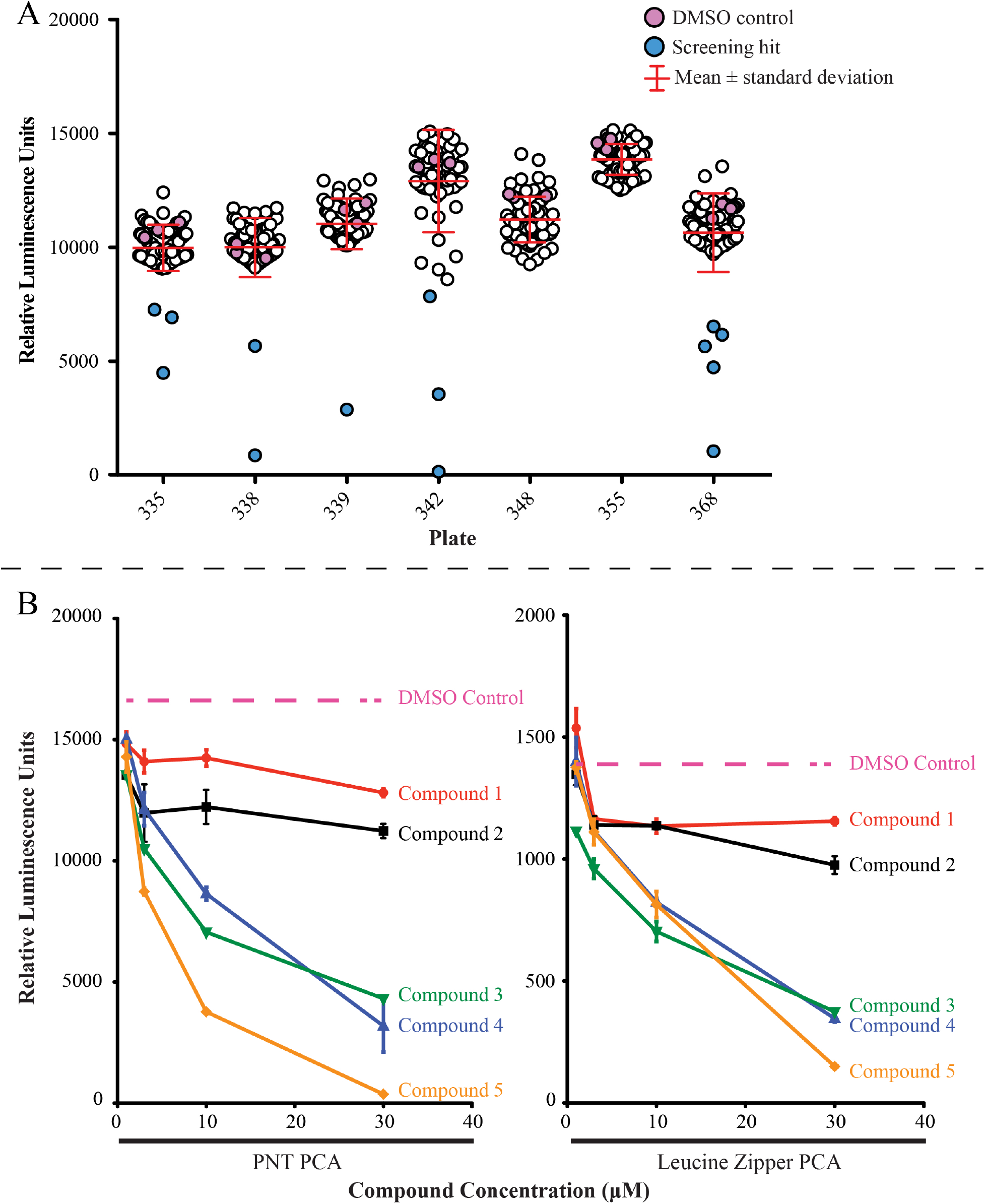
(**A**) Shown are the results of the PNT domain PCA with compound plates from the Prestwick (335, 338, 339), Biomol (342), Sigma (348, 355) and Microsource (368) libraries. The circles represent the luminescence reading of each well of a 96-well plate. The red bars represent the mean relative luminescence and standard deviation of that plate. Pink circles represent wells that were exposed to only DMSO. An initial screening hit is defined as being more than two standard deviations away from the mean (blue circles) (**B**) Compounds identified as hits in the initial high-throughput screen were retested for a concentration-dependent response in the PNT domain and leucine zipper PCAs. Compounds were tested in duplicate at 1, 3, 10 and 30 μM and cells were visually examined to determine toxicity or compound precipitation. Those that were not cytotoxic and elicited a similar response in the both PCA systems, were likely inhibitors of split-luciferase reconstitution, luciferase activity itself or another variable. Compounds 1-2 represent examples of artifacts of the original, large-scale screen whereas compounds 3-5 represent examples of concentration-dependent responses. Compound 1: (±)-pindobind; Compound 2: gitoxigenin diacetate; Compound 3: rosolic acid; Compound 4: gambogic acid amide; Compound 5: 3,4-dimethoxydalbergione.

Secondary testing of screening hits was performed in the PNT domain PCA and the control leucine zipper PCA systems (Fig. 3B). Compounds were re-tested in duplicates at 1, 3, 10 and 30 μM, and cells were examined under the microscope for any toxic effects or evidence of compound precipitation. Out of 179 compounds retested, 83 did not decrease the luminescence in a concentration-dependent manner and were likely artifacts of the initial screens. All 96 of the remaining compounds that showed a concentration-dependent decrease of luminescence in the PNT domain PCA exhibited the same pattern with the leucine zipper PCA. Thus, changes in luminescence were likely due to factors other than inhibition of the PNT domain association.

In addition to the secondary cellular screening, 13 of the 96 compounds were purchased and tested for binding *in vitro* to the PNT domain using NMR spectroscopy. The ^15^N-HSQC spectra of ^15^N-labelled A93D-ETV6^43-125^ was recorded upon progressive titration with each compound. Amide ^1^H^N^ and ^15^N chemical shifts are highly sensitive to even subtle structural changes accompanying ligand binding.^37^ In no case were any chemical shift perturbations observed, indicating no detectable binding (Supplemental Table S3).

### Development of a Yeast Two-hybrid Assay to Screen for Inhibitors of PNT Domain Association

The yeast two-hybrid assay involved several plasmids in auxotrophic yeast strains (Fig. 1B). The bait plasmid pIS586 encoded a fusion between the *POU* DNA-binding domain and the A93D-PNT domain, the prey pIS591 encoded a fusion between the NLS-B42 activation domain and the V112E-PNT domain, and the prey control pIS580 lacked the PNT domain. To validate the assay, we showed that the yeast strain ISY361+/+ containing the “bait + prey” plasmids indeed grew in the absence of His in the medium, whereas the “bait + control” strain ISY361+/− lacking the V112E-PNT domain showed little to no growth (Fig. 2B). Growth was restored upon addition of His to the latter, indicating that lack of growth was specifically due to lack of *HIS3* gene expression.

### High-throughput Screening using the Yeast Two-hybrid Assay

In total, over 8,000 compounds were screened in the yeast two-hybrid assay. The effects of the compounds on yeast cell growth were displayed as histograms for each library, with representative examples in Fig. 4A. In general, the libraries showed a narrow growth distribution range, with most compounds affecting growth by less than 10%.

**Figure 4.**
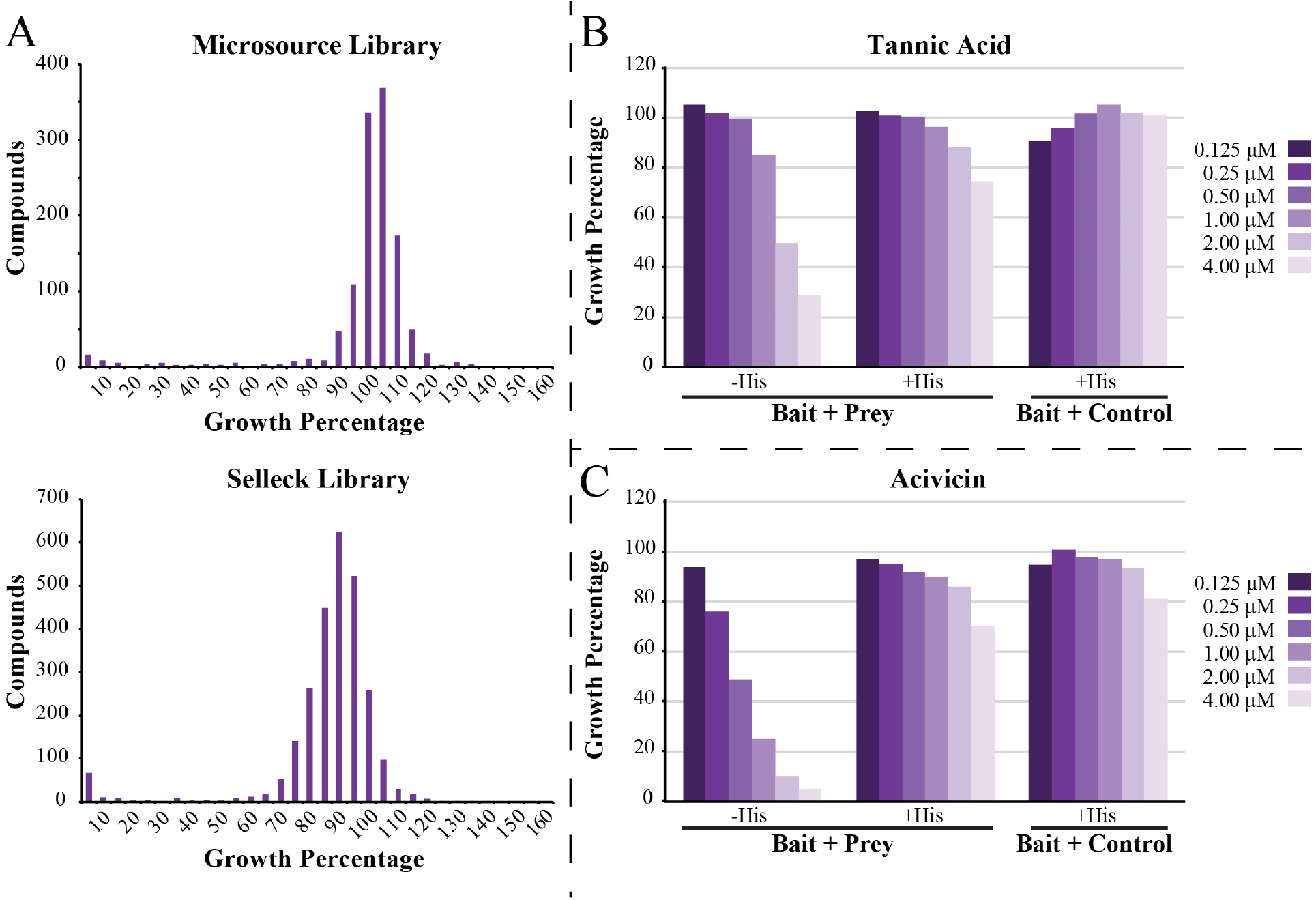
(**A**) Shown are two compound libraries that were tested in the yeast two-hybrid assay with ISY361+/+ co-expressing bait (A93D-PNT Domain/POU DBD) and prey (V112E-PNT domain/AD) plasmids. The histograms summarize the number of compounds versus growth of untreated cells, defined as 100%. Compounds causing reduced growth were subsequently carried on to secondary testing. Both tannic acid (**B**) and acivicin (**C**) showed concentration-dependent inhibition of the growth of yeast containing the “bait + prey” plasmids. This could be rescued through addition of His, indicating that neither compound is toxic to the cells. A similar pattern was seen with ISY361+/− yeast harboring the “bait + control” plasmids, whereby His rescues the cell growth in absence of a functioning *HIS3* gene. Acivicin was later found to be an inhibitor of γ-glutamyltransferase which is necessary for histidine synthesis.

Secondary screening was carried out using multiple concentrations of 242 of the hits from the primary screen. Several compounds, such as tannic acid (Fig. 4B) and acivicin (Fig. 4C), showed a decrease of cell growth in the “bait + prey” strain that could be restored with addition of histidine and exhibited no decreased cell growth in the “bait + control” strain. It was subsequently recognized that acivicin is a glutamine analogue and can inhibit γ-glutamyltransferase,^38^ which is needed for histidine biosynthesis. While such compounds are false positives of the two-hybrid screen, they do validate that a growth response should be observed if a compound interferes with histidine biosynthesis through inhibition of PNT domain association.

In total, three compounds (paromomycin, tannic acid, sanguinarine) passed the secondary screening and have no reported impact on the biosynthesis of histidine. Thus, they were purchased for final testing utilizing NMR spectroscopy. NMR-monitored titrations were carried out with both ^15^N-labelled A93D- and V112E-ETV6^43-125^. In all cases, no amide ^1^N^H^-^15^N chemical shift perturbations we observed, indicating that the three compounds do not detectably bind to the monomeric PNT domains *in vitro* (Supplemental Table S3). The reasons underlying their effects on the yeast two-hybrid screen are currently unknown.

### Virtual Screening for Inhibitors of ETV6 PNT Domain Self-Association

Virtual screening for ligands predicted to bind the ETV6 PNT domain self-association interfaces was performed using the BUDE algorithm with a high-performance computer cluster. Over 160 million total poses of ~ 8 million ligands from the ZINC8 library, averaging 20 conformers per molecule, were tested. Virtual docking of these molecules was initially directed towards 15 Å × 15 Å × 15 Å search boxes centered on the A93 and V112 interfacial regions of the PNT domain (Fig. 5).

**Figure 5.**
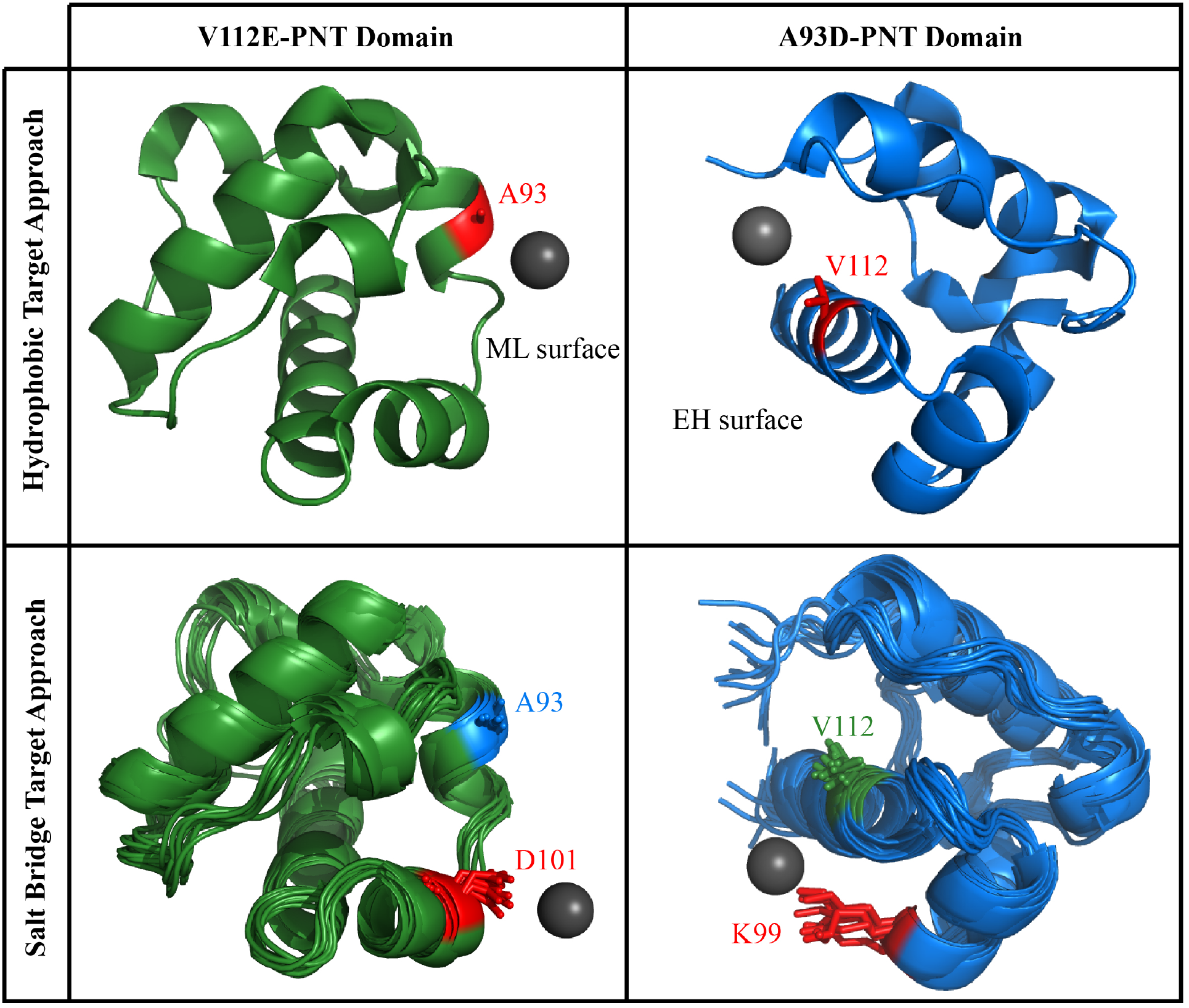
General overview of the BUDE screening. The origin for the starting position of the ligand in the virtual screen is indicated by the grey sphere and the search space was centered around this point. To target the hydrophobic interfaces (upper), PNT domain structures were taken from (PDB: 1lKY). To target the flanking salt bridge (lower), 10 structures from a 100 ns molecular dynamics simulation were chosen.

After running BUDE targeted to each PNT domain hydrophobic interface, the top 500 compounds predicted to have the best binding energies were selected for short MD simulations. The simulations were carried out to determine if the compounds would theoretically remain associated with the PNT domain over a 10 ns sampling. The coordinate RMSDs of each ligand throughout the simulation versus its initial, midpoint (5 ns) and endpoint (10 ns) pose were plotted. Based on these RMSD plots, ligands were scored as “excellent”, “good”, “mediocre” and “bad” (Fig. 6). In parallel, every compound in the top 500 was visually analyzed for common structural motifs and suitable interactions.

**Figure 6.**
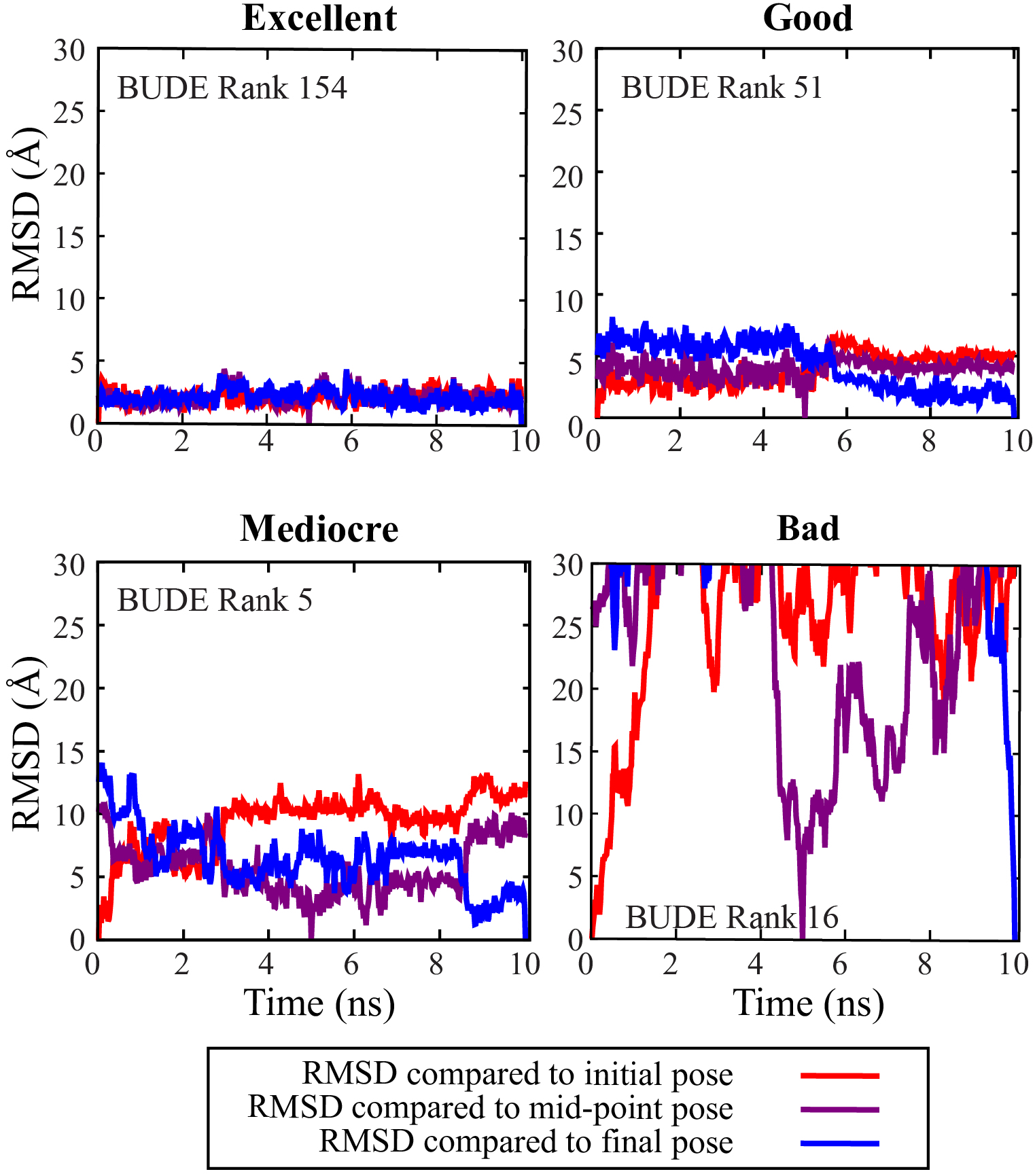
Coordinates of selected ligands over the course of 10 ns MD simulations were compared to the starting (red), midpoint (purple) and endpoint (blue). Ligands were deemed as “excellent” (low RMSDs), “good” (RMSDs < 10 Å), “mediocre” (RMSDs > 10 Å) or “bad” (RMSDs > 30 Å, indicating dissociation).

Compounds selected for testing by NMR spectroscopy and the screening assays had excellent or good scores in MD simulations, appeared to interact closely with the PNT domain, were commercially available and represented a range of chemical motifs to increase diversity. Preference was given to compounds that were predicted to bind in multiple conformers. Several additional compounds that had a BUDE ranking > 500 were also selected due to possessing certain structural motifs, such as carboxylates or protonated amines, that could potentially interact with complementary motifs of the PNT domain. Collectively, this resulted in 35 and 17 compounds predicted to bind the hydrophobic interfaces centered around A93 (ML surface) and V112 (EH surface) of the V112E- or A93D-PNT domain monomers, respectively (Supplemental Fig. S1).

A second screening was subsequently carried out to target the interfaces around K99 and D101. These residues form an intermolecular salt bridge (Fig. 5). Furthermore, to help account for protein dynamics, an ensemble of protein structures taken at 10 ns steps of 100 ns MD simulations were used for docking. To accelerate the process, we also focused on only the top 200,000 compounds from the first BUDE screen and their respective conformers. In total, ~ 4 million poses of ligands were tested for docking to the 10 different A93D- or V112E-PNT domain structures. The resulting top 500 candidates were visually inspected and 10 compounds with chemical diversity were selected for testing (Supplemental Fig. S3).

### Experimental Testing of the Virtual Screening

A total of 60 candidate compounds from two BUDE screens were purchased for further testing. All compounds were used for NMR-monitored titrations with samples of ^15^N-labelled A93D- or V112E-ETV6^43-125^, as appropriate. None were found to bind with any detectable affinity as evidenced by the lack of amide ^1^N^H^-^15^N chemical shift perturbations (Fig. 7A and Supplemental Table S3). In parallel, the compounds were screened in both the yeast two-hybrid and the mammalian cell assays. No compounds were growth inhibitory in the yeast assay, and in the PNT domain PCA, one compound was identified as a screening hit (Fig. 7B). However, this compound also demonstrated reduced luminescence in the leucine zipper PCA, possibly inhibiting luciferase or other confounding factors related to cellular proliferation. Despite extensive efforts, virtual screening did not lead to the identification of any compounds that bind *in vitro* to the ETV6 PNT domain.

**Figure 7.**
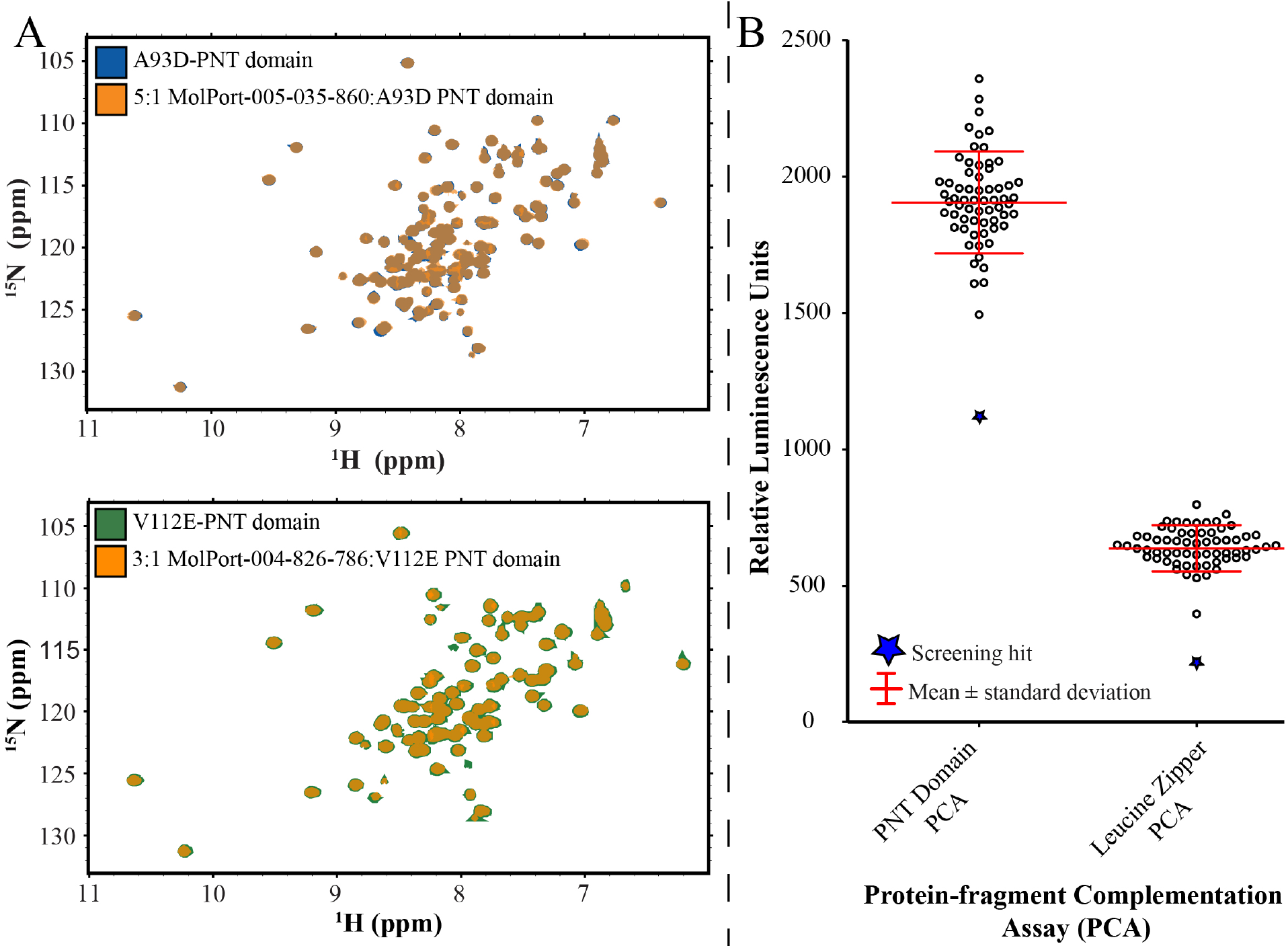
(**A**) Shown are overlaid ^15^N-HSQC spectra of the ^15^N-labelled A93D-ETV6^43-125^ in the absence (blue) and presence of a 5:1 molar ratio (orange) of a BUDE screening hit MolPort-005-035-860 (top), and the ^15^N-labelled V112E-ETV6^43-125^ in the absence (green) and presence of a 3:1 molar ratio (orange) of a BUDE screening hit MolPort-004-826-786 (bottom). The lack of any amide chemical shift perturbations indicated no detectable binding. (**B**) Luminescence readings of 60 compounds, identified through the BUDE virtual screens, tested with the mammalian cell split luciferase assay. One compound (MolPort-005-970-014; blue star), reduced luminescence in both the PNT domain and leucine zipper PCAs. The lower signals relative to previously presented studies is due to the use of a BioTek Neo 2 plate reader.

## Discussion

In this study, we established three complementary approaches to screen for potential inhibitors of ETV6 PNT domain polymerization. Unfortunately, we did not succeed in discovering any small molecules that bound the PNT domain *in vitro* or disrupted its self-association *in vivo*. Nevertheless, valuable insights were gained that should facilitate future studies aimed at modulating protein-protein interactions with systems similar to ETV6.

The mammalian and yeast cell assays were designed for unbiased high-throughput screening of compounds in a cellular context, with a readout that is dependent upon disruption of a heterodimer model of the ETV6 polymer. However, in the PCA assay, screening hits can arise not only from inhibition of heterodimerization, but also from perturbation of split luciferase reconstitution or luciferase activity, as well as due to cytotoxicity or inhibition of cellular proliferation. Therefore, the leucine zipper PCA was critical as a control assay to determine whether any reduction in luminescence was due to the latter confounding effects. Similarly, in the yeast screening assay, the use of a control vector and addition of histidine to secondary testing of screening hits for growth rescue were used to eliminate any compounds that caused cell death or inhibition of the histidine biosynthetic pathway.

It is worth emphasizing that the ETV6 A93D- and V112E-PNT domains heterodimerize with low nM affinity.^18,19^ This certainly presents a challenge in discovering potentially rare compounds in chemical libraries with comparable affinities to effectively disrupt this tight protein-protein interaction. A mitigating strategy might involve the initial use of PNT domain variants with additional mutations to weaken their association and thereby reduce the stringency of the assay. Such variants may be inspired by detailed biophysical analyses of the ETV6 PNT domain interfaces.^39^ In addition, it would be beneficial to use cell-based “up” assays that positively select for disruption of PNT domain association or employ a readout that increases, rather than decreases, with this result.

To this end, we explored use of a yeast two-hybrid assay where expression of *HIS3* and *LacZ* reporter genes are dependent upon PNT domain heterodimerization. However, this assay as implemented was not useful for discovery of inhibitors because potential hits in small molecule screens would prevent growth on medium lacking His, and loss of *LacZ* expression, which could also be produced by various alternative effects unrelated to inhibition of the PNT domain interaction. Ideally, the screening assay could be modified to include a reporter such as *CAN1* or *URA3*, where inhibitors of the interaction would prevent activation, allowing detection by growth on canavanine or 5-fluoroorotic acid, respectively.^40^ Alternatively, yeast-based strategies could also be used such as the repressed transactivator system, where interaction of bait and prey fusions cause repression of gene expression, and inhibitors of the interaction cause gene activation, which is detected by growth of yeast on selective media.^41^ However, this system is currently limited to use with Gal4 DNA-binding domain fusions, and we were unable to detect interaction of PNT fusions in yeast using the Gal4 DNA-binding domain linked to the PNT domain.

In parallel to cellular assays, we undertook a targeted virtual screening approach to search for compounds that bind the known polymerization interfaces of the ETV6 PNT domain. Although BUDE has been successful in identifying inhibitors of PPIs,^26^ and numerous potential ligands were predicted to bind the ETV6 PNT domain *in silico*, none were found to do so when tested *in vitro* with NMR spectroscopy or *in vivo* with cellular assays. However, common motifs that were enriched may serve as starting structures for rational design or further high-throughput screening experiments.

We recognize that this target is particularly challenging since the ETV6 PNT domain polymerizes with high affinity through two relatively large flat interfaces. These comprise complementary central hydrophobic patches ringed by hydrophilic and charged residues with relatively flexible sidechains. Tightly bound ligands typically occupy depressions or cavities on the protein surface, but these features are absent with the PNT domain. However, in the case of targeting the intermolecular salt-bridge, the virtual ligands seemed to be predicted to preferentially bind to certain PNT domain structures taken from short MD simulations. This flexibility of the protein may expose conformations better suited for small molecule interactions.

It is unfortunate that some of the best hits from the virtual screen were not commercially available. Experimental testing of these compounds, as well as more *in silico* hits, might yield success. Indeed, a “similarity paradox” has been described where minor chemical modifications of otherwise similar molecules can render them active or inactive.^42^ This paradox may suggest that, in our efforts to test a diverse set of compounds that sampled chemical space, we missed a high-ranking compound with affinity for the ETV6 PNT domain. Such a large-scale screening approach may be effective, as many protein-protein interaction inhibitors are found through traditional cellular screening only after screening in excess of hundreds of thousands of compounds and carrying out structure-activity relationship studies to optimize initially detected weak-binding leads.^43^

Fragment-based drug design and disulfide tethering, combined with combinatorial chemistry, are two approaches that could also be undertaken as the next steps in developing an inhibitor against PNT domain polymerization.^44–46^ Inspiration can also be found in the design of cyclic peptides and helix peptide mimetics that weakly bind the SAM domains of the Ship2 and EphA2 receptors.^47,48^ In all cases, virtual screening could be used to narrow down potential chemical motifs of interest for a more targeted screening approach. Collectively, such a multi-pronged strategy may lead to the discovery of a potentially new class of therapeutics that prevent the polymerization of chimeric oncoproteins resulting from a wide array of *ETV6* chromosomal translocations.

## Acknowledgements

We thank Mark Okon for assistance with NMR spectroscopy, Stephen Michnick for providing starting plasmids for the split luciferase PCA, Andrew Wilson for the PoPPI library, and Poul Sorensen for advice on ETV6 translocations.

Supplemental material is available online with this article.

## Supplemental Material

**Supplemental Table S1.**
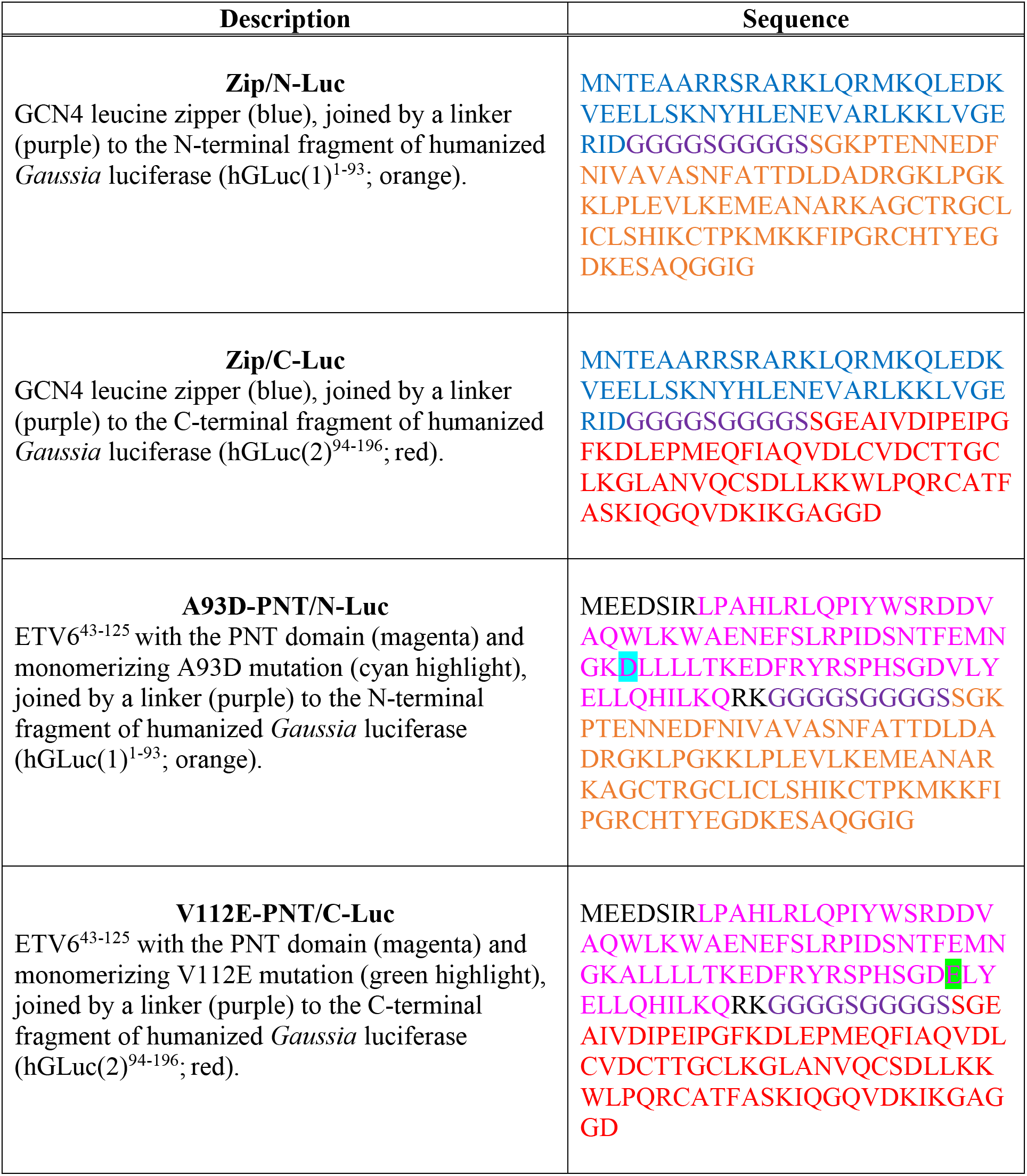
Protein sequences of selected PCA constructs

**Supplemental Table S2.** Luminescence readings of the PNT domain PCA from transiently transfected cells and from stable pooled transformants.

| **Plasmid(s) Transfected or Conditions** | **Hours Post-Transfection** | **Luminescence Reading** |
| --- | --- | --- |
| DMEM, no HEK293 cells | 48 | 20 |
| HEK293 cells with a mock (no DNA) transfection | 48 | 25 |
| V112E-PNT/C-Luc | 48 | 670 |
| V112E-PNT/C-Luc | 72 | 500 |
| A93D-PNT/C-Luc | 48 | 450 |
| A93D-PNT/N-Luc | 48 | 690 |
| A93D-PNT/N-Luc and V112E-PNT/N-Luc | 48 | 70 |
| A93D-PNT/N-Luc and A93D-PNT/C-Luc | 48 | 2500 |
| A93D-PNT/C-Luc and V112E-PNT/N-Luc | 48 | 320,000 |
| A93D-PNT/C-Luc and V112E-PNT/N-Luc | 72 | 165,000 |
| A93D-PNT/N-Luc and V112E-PNT/C-Luc | 48 | 91,000 |
| A93D-PNT/N-Luc and V112E-PNT/C-Luc | 72 | 68,000 |
| Zip/N-Luc and Zip/C-Luc<sup>1</sup> | 48 | 45,000 |
| Zip/N-Luc and Zip/C-Luc<sup>1</sup> | 72 | 22,000 |
| Stably expressing PNT domain PCA | N/A | 14,000 |
| Stably expressing leucine zipper PCA | N/A | 1,020 |

<sup>1</sup>Denoted as Zip-hGLuc(1) and Zip-hGLuc(2) in Remy, I. and Michnick, S.W. *Nat. Methods* **2006**, 3 (12), 977–979.

**Supplemental Table S3.**
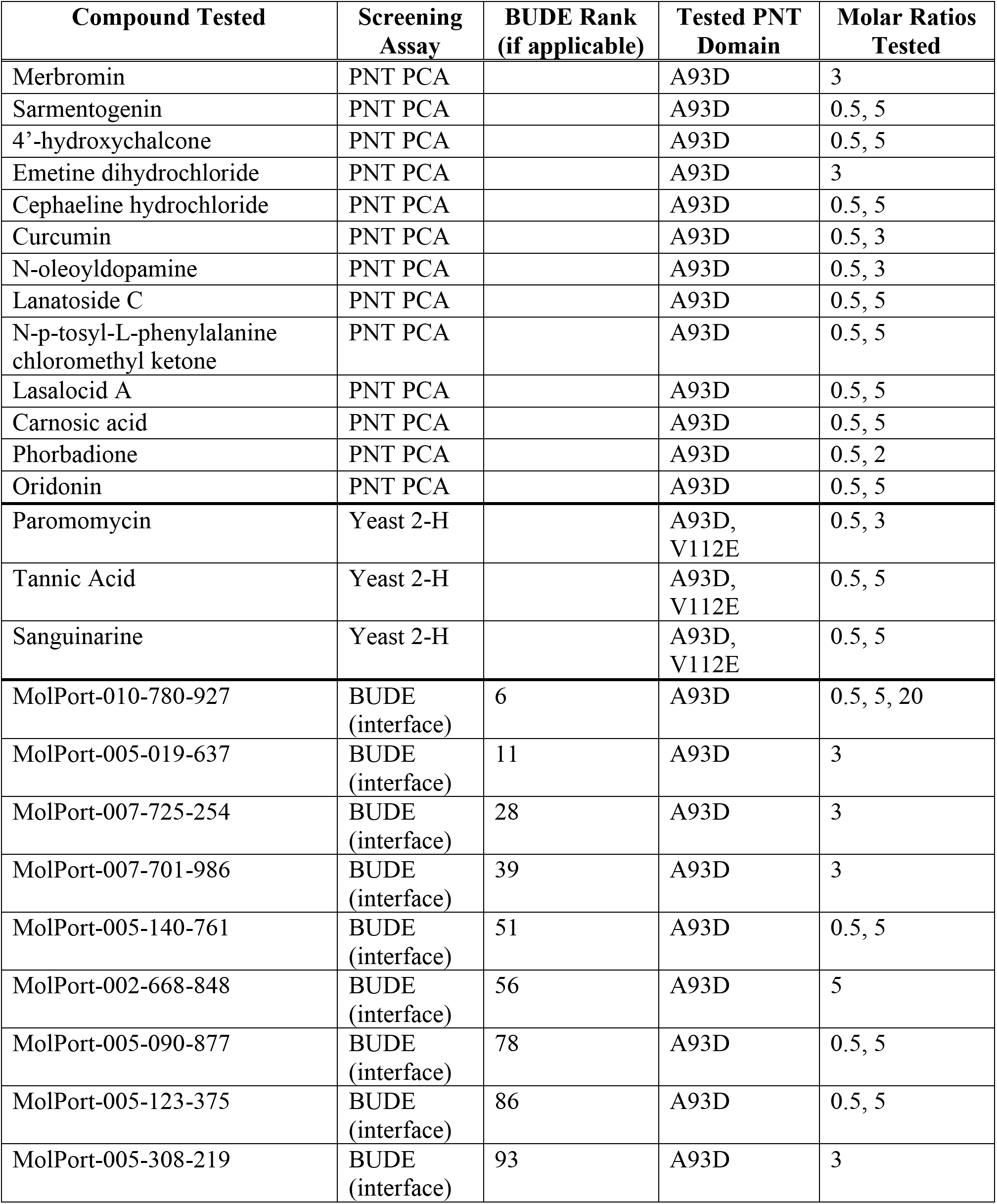
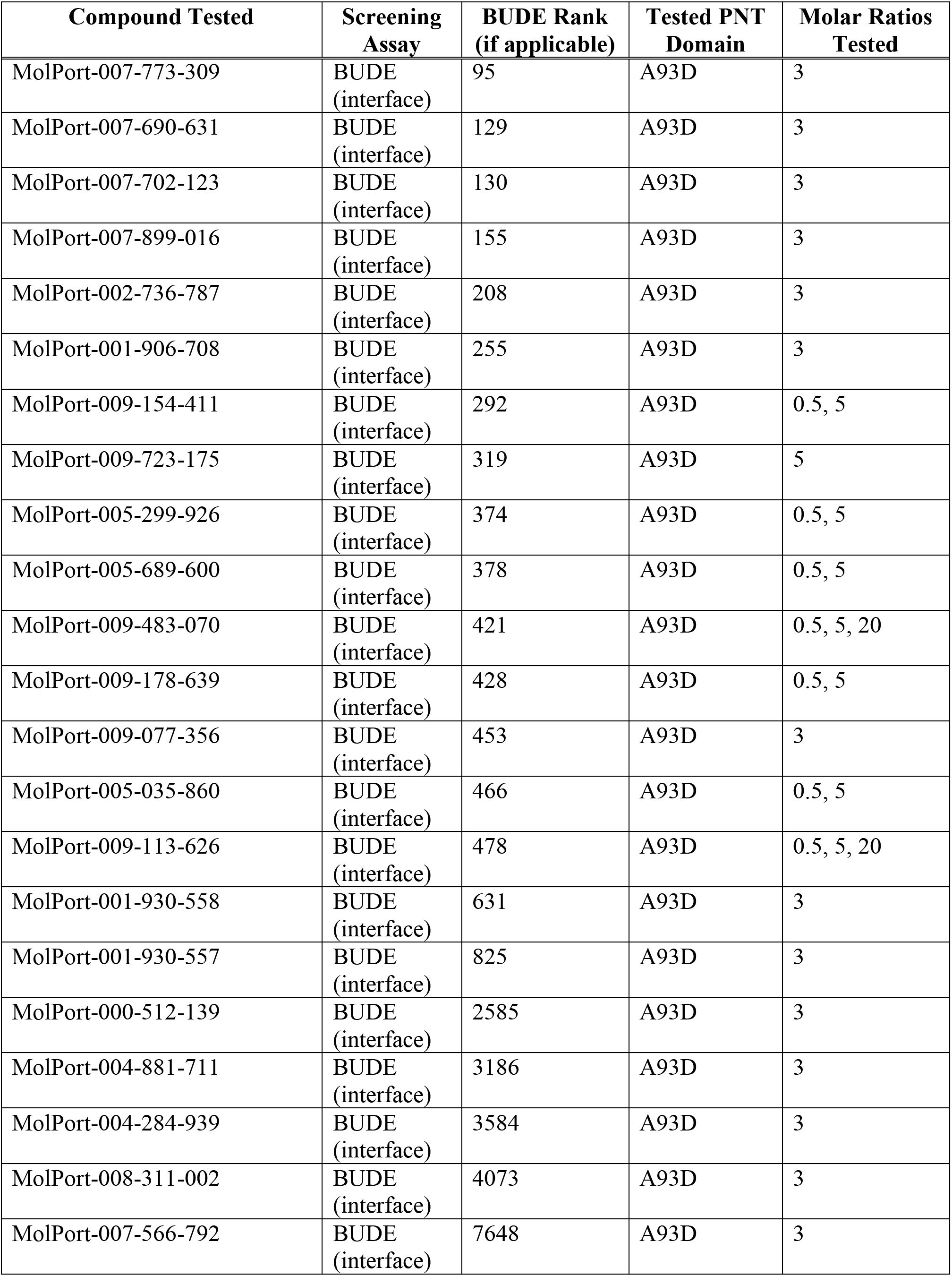
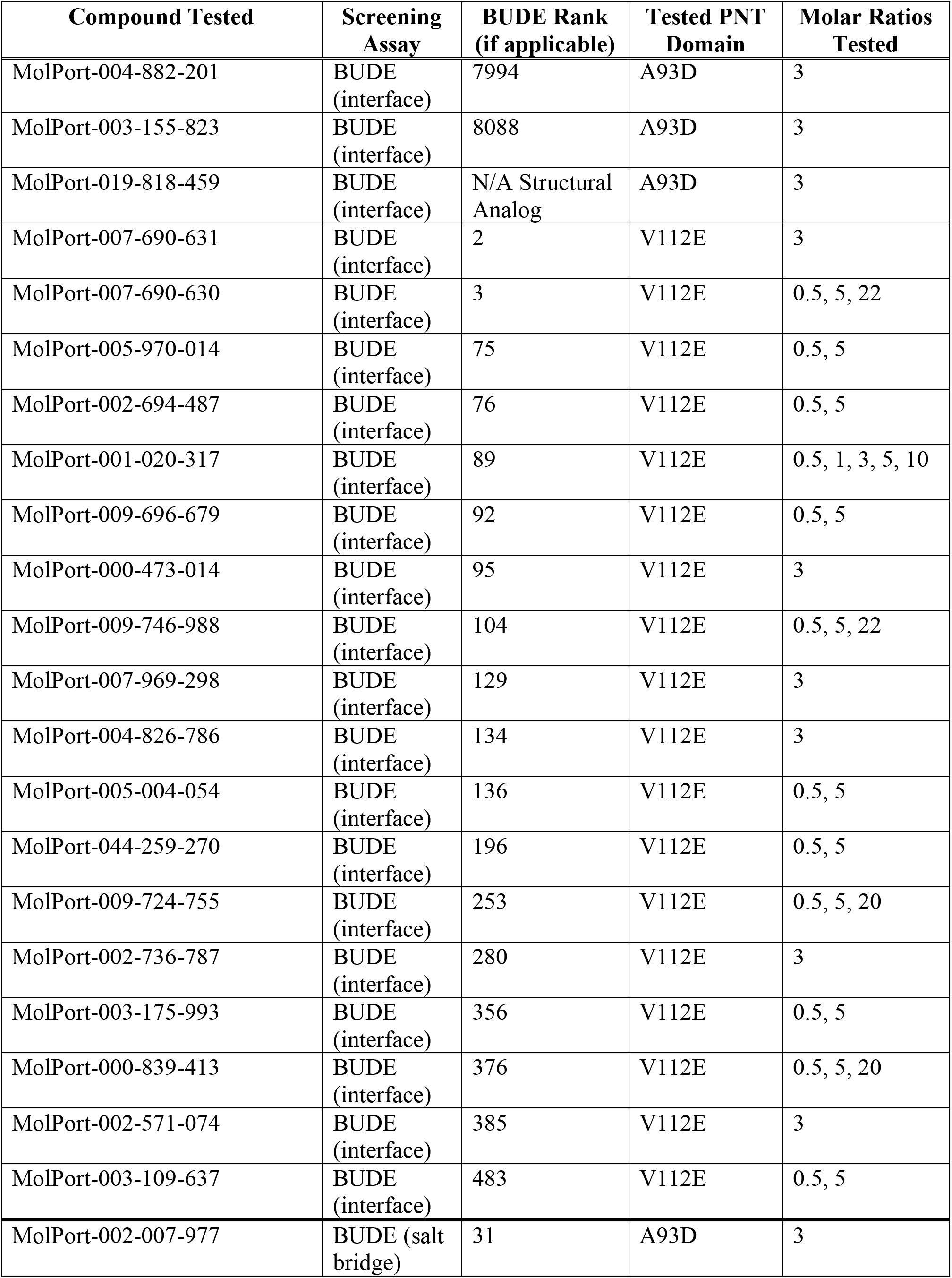
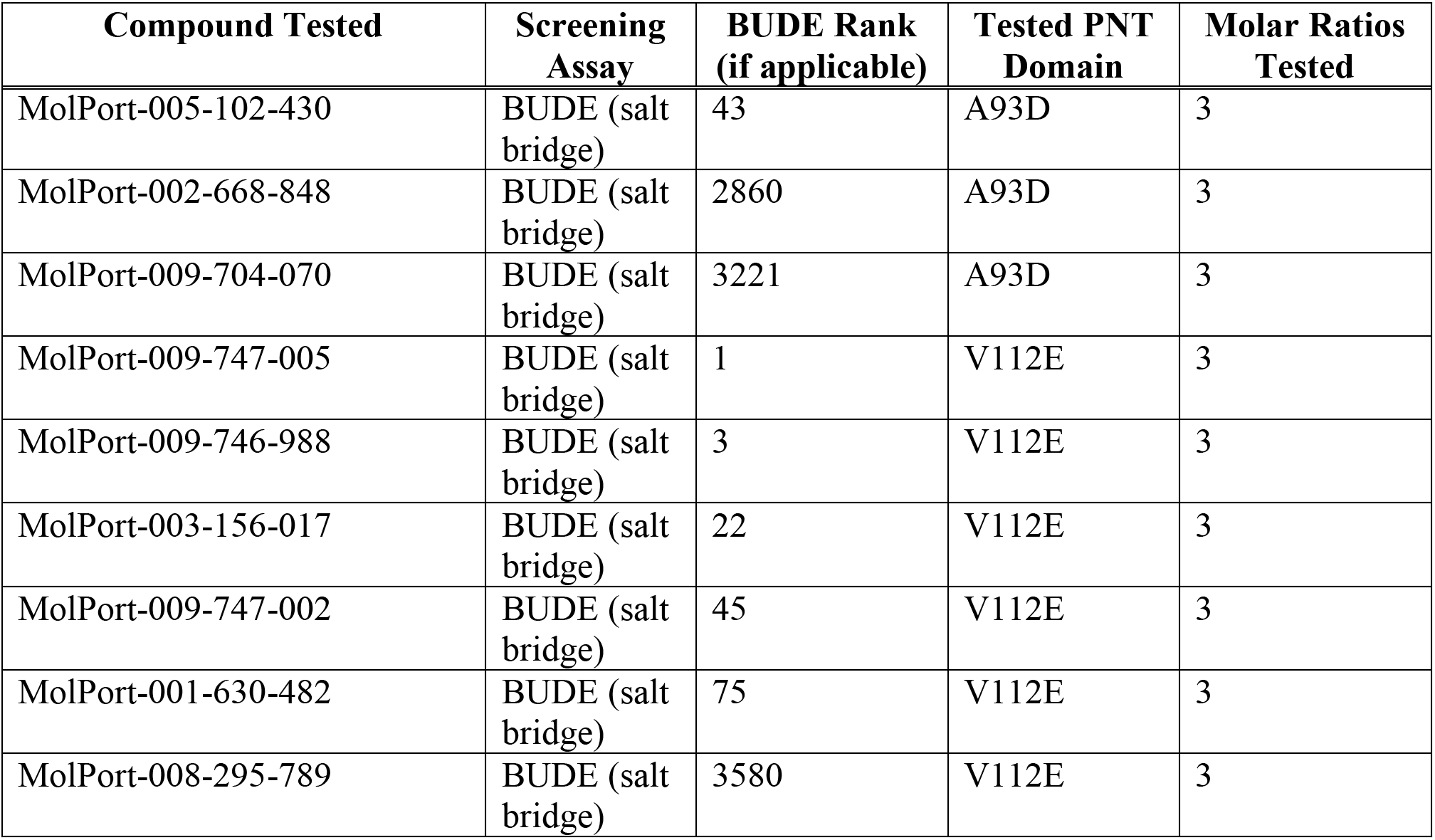
Purchased compounds tested for binding to the A93D- or V112E-PNT domain (in context of ETV6^43-125^) NMR spectroscopy. None bound as evidenced by the lack of any detectable amide ^1^H^N^-^15^N chemical shift perturbations.

**Supplemental Figure S1.**
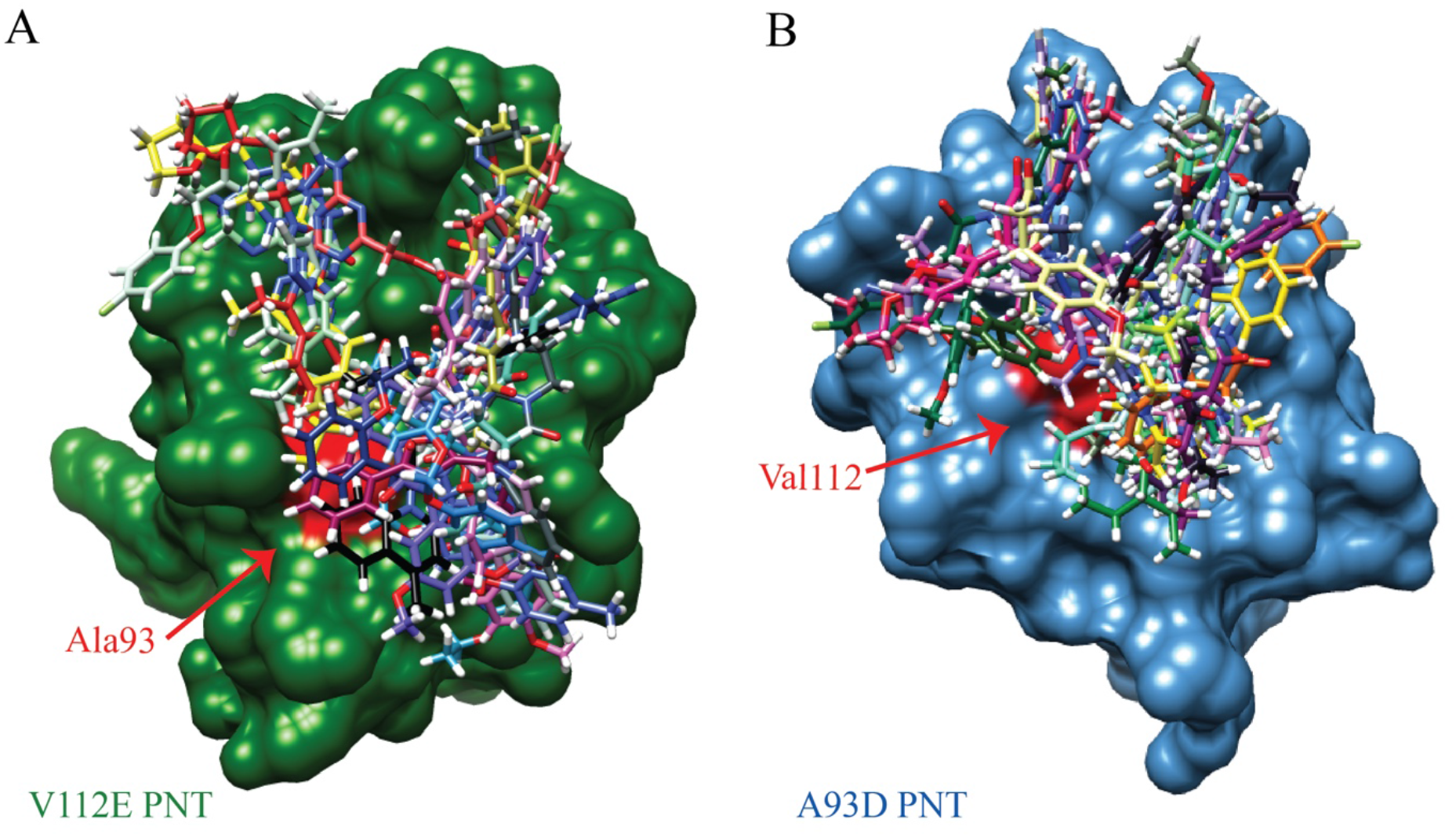
The BUDE compounds tested for binding at the interface and their predicted binding site. (**A**) Rendition of the V112E-PNT domain with compounds predicted to bind the ML surface centered around A93 (red). (**B**) Rendition of the A93D-PNT domain with compounds predicted to bind the EH surface centered around V112 (red). A range of binding locations can be seen. The protein structures were adapted from PDB 1LKY. The superimposed docked compounds are shown in stick format (hydrogen, white; nitrogen, blue; oxygen, red; sulfur, yellow; halogens, neon green; carbon, remaining colors).

**Supplemental Figure S2.**
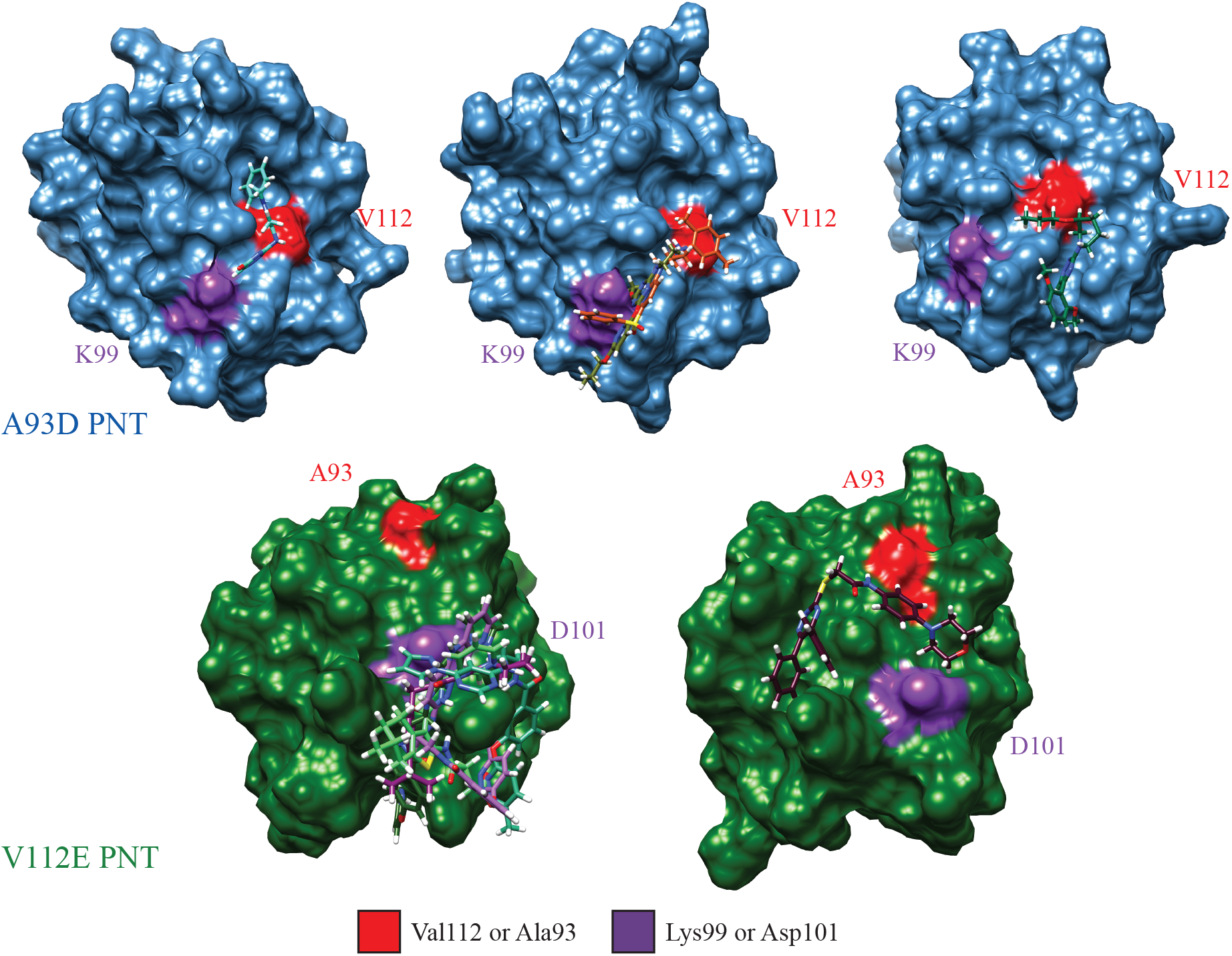
Virtual docking of compounds targeted to the K99-D101 interfacial regions of the PNT domain. Shown are surface models of three A93D PNT domain (blue) and two V112E PNT domain (green) structures, generated by MD simulations, with four and six docked compounds, respectively, that were experimentally tested for binding by NMR spectroscopy. The superimposed docked compounds are shown in stick format (hydrogen, white; nitrogen, blue; oxygen, red; sulfur, yellow; halogens, neon green; carbon, remaining colors).

